# School yard biodiversity determines short-term recovery of disturbed skin microbiota in children

**DOI:** 10.1101/2021.06.18.448749

**Authors:** Jacob G. Mills, Caitlin A. Selway, Torsten Thomas, Laura S. Weyrich, Andrew J. Lowe

**Affiliations:** School of Biological Sciences, The University of Adelaide, South Australia, Australia; Centre for Marine Science and Innovation, School of Biological, Environmental and Earth Sciences, University of New South Wales, Sydney, Australia; Department of Anthropology, The Pennsylvania State University, University Park, USA

## Abstract

The planting and conservation of biodiverse habitat in urban contexts has been proposed as a public health intervention aimed at reducing the prevalence of non-communicable diseases via microbiome rewilding (Mills et al. 2017; Mills et al. 2019). However, our understanding of the effect of urban biodiversity interventions on the human microbiota remains limited, especially on the skin (Hui et al. 2019; Roslund et al. 2020) and in the context of permanent green spaces (Lehtimäki et al. 2018; Selway et al. 2020). Here, we test the short-term response of experimentally disturbed bacterial communities on the skin of healthy children exposed to different school environments – either a ‘classroom’, a ‘sports field’, or a biodiverse ‘forest’ – to understand how exposures to different types of biodiversity may influence skin microbiota. Children exposed to the ‘forest’ had significantly increased skin microbiota diversity when compared to pre-exposure, an effect that increased over three days suggesting long-term effects. The microbiota on children exposed to the ‘forest’ had the largest structural and compositional community change compared to children exposed to ‘sports fields’, which in turn was larger than those who remained in ‘classrooms’. Children exposed to ‘sports fields’ and ‘forests’ also acquired new core bacteria after exposure to green spaces, potentially buffering against disturbances to the skin microbiota’s diversity, while individuals who remained in the ‘classroom’ lost microbes throughout the experiment. Overall, we conclude that urban green spaces can have an enriching influence on the diversity of skin microbiota, including core members shared between all children. These findings have important implications for the design and construction of new school yards and public spaces with respect to biodiversity, health, and human microbiota.

## Main

There is increasing awareness that the skin microbiota is mechanistically important for human health, including immune and physiological responses (Byrd et al. 2018; Chen et al. 2018). The microbiota on the skin, and at other sites in the body, develop mostly through environmental factors and acquirement during early life following principles of community ecology and successional theory (Costello et al. 2012; Arrieta et al. 2014; Byrd et al. 2018; Leung et al. 2018). These environmentally acquired microbiota, in particular the co-evolved ‘old-friends’ microbiota (Rook 2013), have the potential to shape life-long health trajectories (Rackaityte & Lynch 2020).

However, there are everyday medical and lifestyle pratices that can disturb skin microbiota and disrupt diversity and composition (e.g., hand-sanitisation; SanMiguel et al. 2018), and contribute to the development and outcomes of skin diseases (Van Rensburg et al. 2015; Byrd et al. 2018). Moreover, exposure to biodiversity and rich environmental sources of microorganisms has become severely limited in modern cities, as humans spend more time indoors under clean or industrial conditions, which is linked to a decrease in microbial diversity within the human body (Sonneberg and Sonneberg, 2020). Overall, these decreases in microbial exposure can have a number of potential health implications (von Hertzen et al. 2011).

We recently proposed the ‘microbiome rewilding hypothesis’, which suggests that exposures to highly biodiverse urban environments may provide a means to increase microbial diversity in the human body (Mills et al. 2017). Recent experimental research has shown that children in nature-based day-care facilities show increases in bacterial diversity on their skin and in immune function relative to children in conventional or city-based day-care centres (Lehtimäki et al. 2018; Roslund et al. 2020). This provides evidence that simple urban biodiversity interventions could constitute a positive health intervention. However, this has not yet been examined in connection to outdoor environments and outside play, which is critial for our understanding of the mechanistic interactions between a healthy skin microbiota and the broader environment. Specifically, we need to understand the effect of urban green spaces and relative biodiversity quality (e.g., the functional diversity of ecological communities) on the microbial communities, including those of the skin. We hypothesise that exposure to green space will increase diversity and change the composition of the skin’s microbial communities more than staying inside, and that exposure to a more biodiverse green space would have an even greater effect than a less biodiverse area.

To test our hypotheses, we first assigned three participating classes of 10-11 year-old students from a single primary school to one of three different treatment environments of varying biodiversity quality within the school grounds: 1) an indoor ‘classroom’ control (*n* = 20, *n* females = 10), 2) a low vegetation complexity ‘sports field’ (*n* = 14, *n* females = 7), or 3) a high vegetation complexity ‘forest’ (*n* = 23, *n* females = 8). We swabbed the left volar wrists of each group of students before environmental exposure. These swabs gave us pre-exposure information on their skin’s bacterial communities and also acted as an experimental disturbance (i.e., swabbing is an act of cleaning or removing a fraction of the microbiota). We then exposed the students to their assigned environment for 45 minutes – a standard time period spent on individual academic activities in a school setting in Australia. Following environmental exposure, we re-swabbed the same patch of skin to see how environmental exposure had affected the bacterial communities. We repeated this regime at the same time for three consecutive days, always swabbing the same patch of skin. Based on amplicon sequence variants (ASVs) of the bacterial V3-V4 16S rRNA marker gene (Table S1), we performed ecological analyses on whole and core bacterial communities of the skin.

We first tested pre-existing differences between the groups, given that each group of students had been together in their respective classrooms prior to this study for approximately nine months of the 2019 school year (Leung et al. 2018). There was no statistical support for differences in the alpha diversity of the bacterial communities between treatment groups prior to exposure on any of the sampling days (Table S2). However, we did find statistically significant differences in the bacterial community structure between the ‘classroom’ group when compared to the the ‘sports field’ and ‘forest’ groups prior to environmental exposure (weighted-UniFrac, Figure 1b & 1c, Table S4), and we found all three treatment groups were significantly different in their pre-exposure community composition (unweighted UniFrac; Figure 1d & 1e, Table S4). Further, the class assigned to the ‘forest’ exposure treatment had a significantly less diverse core community (i.e., ASVs present in at least 50 % of samples within a group) prior to exposure on days one and three when compared to equivalent samples of the ‘classroom’ group (Figure 3a, Table S3). The structure of the core skin microbiota of the ‘classroom’ group was also significantly different to the ‘sports field’ and ‘forest’ groups (Figure 3b & 3c, Table S5), and each group’s core bacterial composition was significantly different prior to exposure (Figure 3d & 3e, Table S5). These results show that for some community measurements there were unique signatures for the skin microbiota of each group prior to environmental exposure during this experiment. The group uniqueness is consistent with another study suggesting that increasing time spent together can normalize skin microbiota of individuals within groups (Leung et al. 2018). Therefore, we focused our analysis on the impact of environmental exposure within each treatment group.

**Figure 1.**
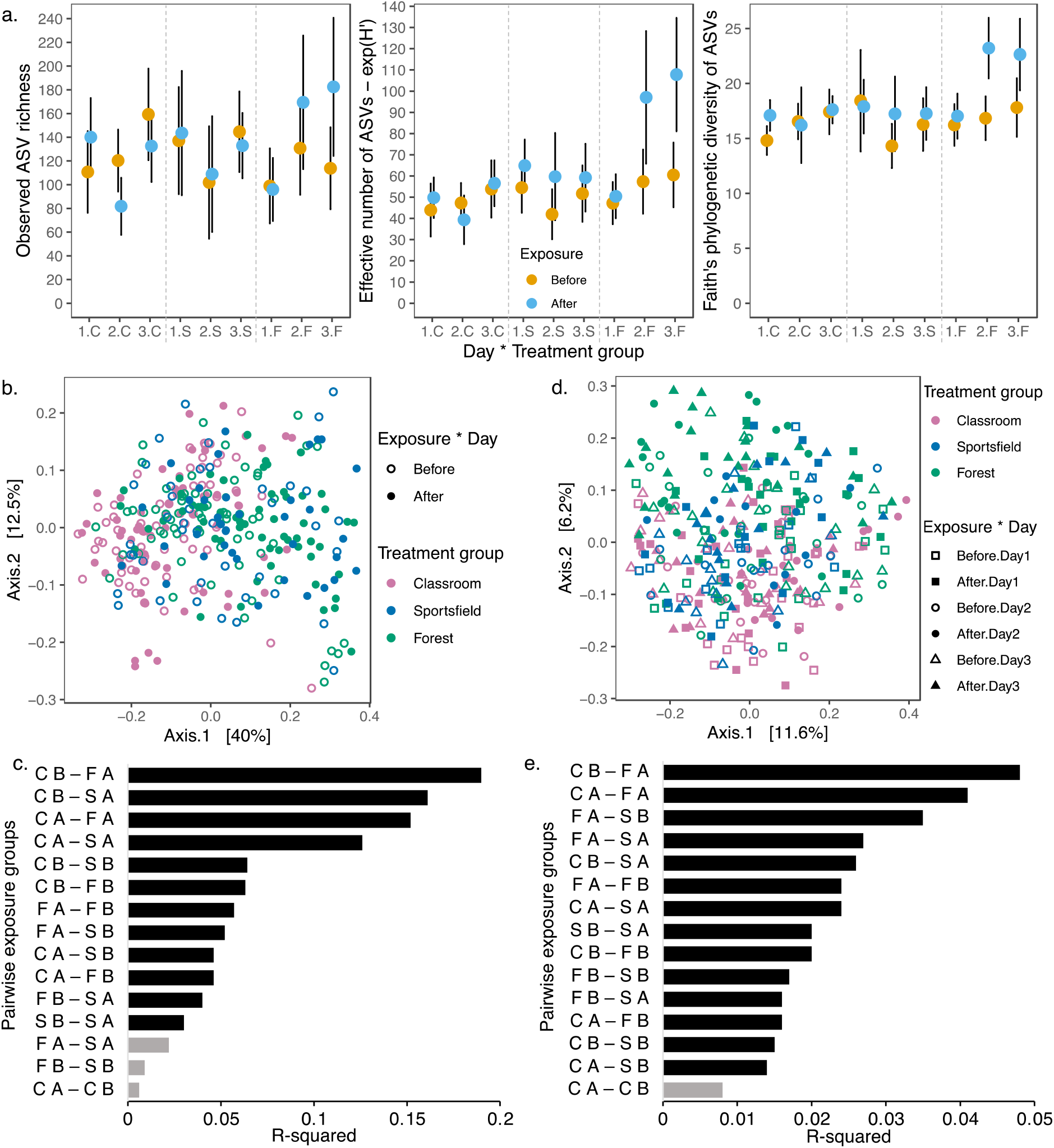
Bacterial ASV communities of children’s wrists ‘before’ and ‘after’ exposure to school yard environments repeatedly sampled across three days. **a**. Observed richness, effective number (exponent of Shannon’s diversity), and Faith’s phylogenetic diversity of ASVs are shown from the wrists of children exposed to three different school yard environments over three consecutive days. Points are means ± 95 % confidence interval. Significantly different pairs are listed in Table S4. ‘1’, day 1; ‘2’, day 2; ‘3’, day 3; ‘C’, classroom; ‘S’, sports field; ‘F’, forest. **b. & d**. PCoA analyses of weighted-UniFrac and unweighted-UniFrac values, respectively, from all skin samples taken ‘before’ and ‘after’ outdoor exposure. Sampling ‘day’ is shown in the unweighted-UniFrac PCoA because it significantly interacted with ‘treatment group’ and ‘exposure’ in the PERMANOVA (Table S4). **c. & e**. Bar plots ranking R^2^ values to show strength of difference between pairs of environmental exposure groups (i.e., classroom group ‘after’ exposure vs. classroom group ‘before’ exposure, CA – CB) from the weighted-UniFrac and unweighted-UniFrac PCoAs, respectively. C, ‘classroom’; F, ‘forest’; S, ‘sports field’; B, ‘before’ exposure; A, ‘after’ exposure. Grey bars represent no significant difference between pairs with α = 0.05, while black bars represent significant difference.

We next investigated the effect that environmental biodiversity quality had on the diversity of the disturbed skin microbiota. The group that spent their class time in the ‘forest’ was the only group to have a significant increase in alpha diversity, as observed richness, effective number, and Faith’s phylogenetic diversity of ASVs were all higher in post-disturbance samples compared to pre-disturbance (Figure 1a, Table S2). This effect also appeared to compound from day one to day three (Figure 1A, Table S2). The school forest is likely a richer source of environmental bacteria than a classroom and sports field, as previous studies have shown that floristically diverse urban green spaces have a richer soil microbiota than less diverse spaces (Hui et al. 2017; Mills et al. 2020). Therefore, we examined the diversity of microbial communities present in each of the these three environments (i.e., swabs from the classroom bench tops (not the student’s tables) and tables (desks), sports field turf, forest soil, and forest leaves). The alpha diversity in our environmental samples from the classroom bench tops were comparable with forest soil and leaves (Figure S1). In this respect, there may be other dynamics, such as indoor and outdoor air-flow differences influencing aerial entrainment of microorganisms or the lack of touch contact to particular surfaces, that can contribute to increases in alpha diversity in disturbed skin microbiota of school age children in some environments relative to others (Robinson et al. 2020).

We next analysed the phylogenetic structure and composition of the bacterial communities of childrens’ skin using weighted and unweighted UniFrac distance matrices. The ‘forest’ and ‘sports field’ treatment groups had statistically significant changes in their bacterial community structure after exposure to their respective environments, while the ‘classroom’ group did not (Figure 1b & 1c, Table S4). Composition of the skin’s bacterial communities had a significant interaction between the factors ‘treatment group’, ‘exposure’, and ‘day’ (Figure 1d & 1e, Table S4), as the composition changed significantly for the ‘forest’ and ‘sports field’ groups after each exposure (Figure 1d & 1e, Table S4). However, the ‘classroom’ group’s skin microbiota became less variable over the repeated days (dispersion P < 0.05, Figure 1d), and this did not occur in the other groups (dispersion P > 0.05).

We explored ASVs shared between human and environmental samples, as these likely represent microorgansims that were transferred onto the skin of children from these exposures. The group that went to the ‘forest’ lost 26.2 % (632 of 2,410) of their ‘before’ exposure ASVs, but gained 1,420 for a total of 3,198 ASVs ‘after’ exposure. For the ‘forest’ group, 171 of the acquired ASVs were also found on forest leaf surfaces, 343 in forest soil, and 342 on either forest leaves or in soil, while 564 had an unknown origin during environmental exposure (Figure 2b). In contrast, the group that went to the ‘sports field’ lost 38.5 % (742 of 1,929) of their ‘before’ exposure ASVs and gained 692 during exposure for a total of 1,889 ASVs after-exposure, with 212 found on the sports field leaves (turf grass) and 480 from an unknown origin (Figure 2c). Further, the ‘classroom’ group lost (33.6 %) 780 of 2,318 (33.6 %) ASVs from their ‘before’ exposure samples but gained only 657 ASVs (totalling 2,195 ASVs ‘after’ exposure) found on bench tops (174), tables (14), either bench tops or tables (11), or unknown origin (458) (Figure 2d). We observed more total ASVs from unknown origin when we split the samples by treatment group (i.e., Figures 2b-d) than when they were pooled (i.e., Figure 2a), suggesting that some ASVs are either moving between these environments, likely through the air, and colonising children in other spaces or that we are not detecting some of the rarer bacteria in all samples. Nevertheless, these results indicate a strong human skin-environment interaction with their respective exposure environment. While significant differences in composition are observed within 45 minutes of exposure, the compounding effect over successive days (Figure 1a) suggests that exposure to more biodiverse areas could have longer term effects on diversifying children’s skin microbiota. Together, these results suggest that biodiversity quality of green space can play a role in the enrichment of skin bacteria over time.

**Figure 2.**
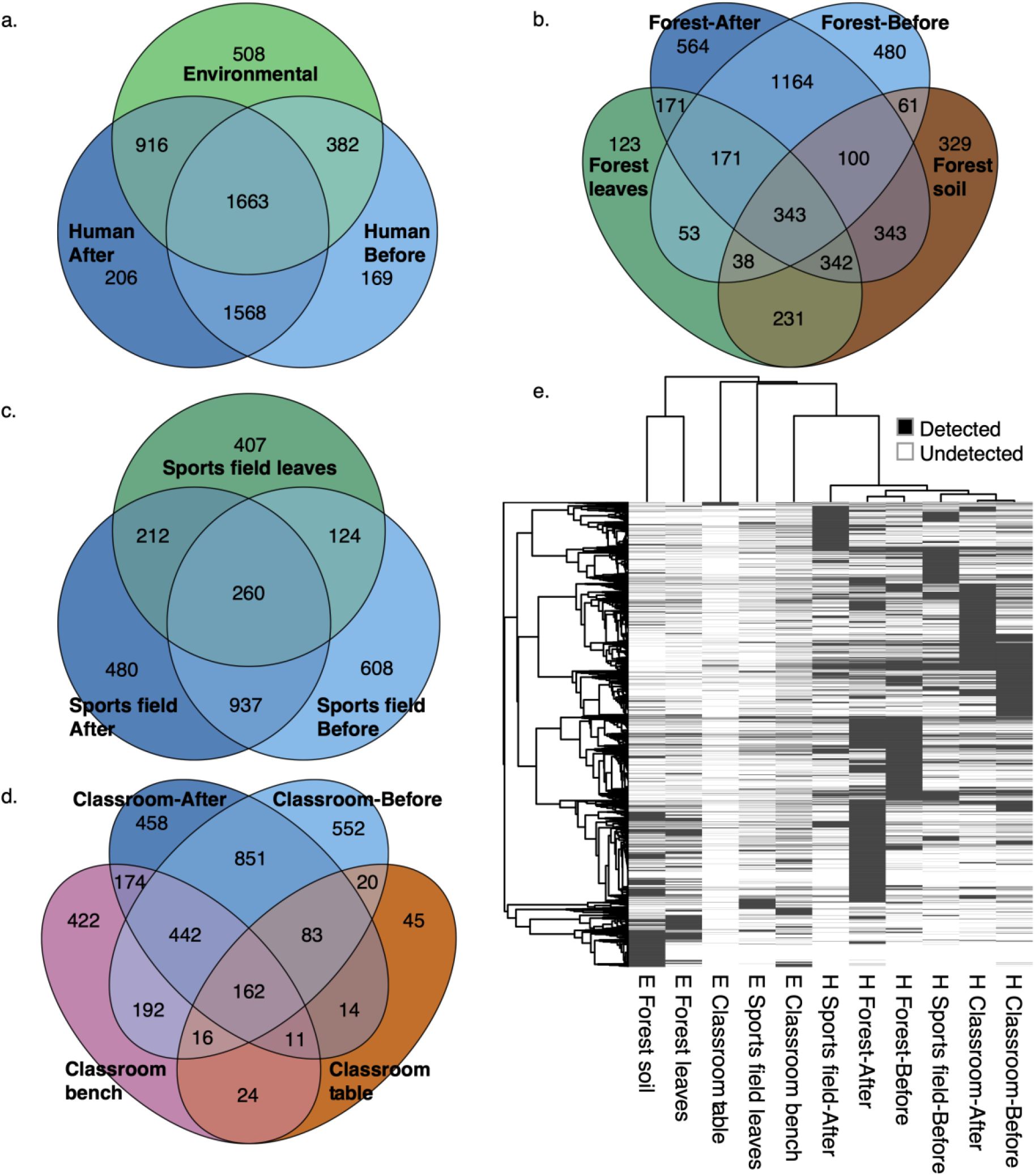
Shared and unshared bacterial community ASVs between human samples and environmental samples. **a**. Total shared and unshared bacterial ASVs between environmental samples and human samples collected ‘before’ and ‘after’ exposure. **b**. Total shared and unshared bacterial ASVs between the ‘forest’ environmental samples (soil and leaf surfaces) and human samples from the ‘forest’ treatment group collected ‘before’ and ‘after’ exposure. **c**. Total shared and unshared bacterial ASVs between the ‘sports field’ environmental samples (leaf surfaces) and human samples from the ‘sports field’ treatment group collected ‘before’ and ‘after’ exposure. **d**. Total shared and unshared bacterial ASVs between the ‘classroom’ environmental samples (bench tops and work tables) and human samples from the ‘classroom’ treatment group collected ‘before’ and ‘after’ exposure. **e**. Heatmap of detected community bacterial ASVs by sample type with clustering representing Pearson correlation between columns (samples) and between rows (ASVs). H and E on the x-axis represent human and environmental sample types, respectively.

Core microbiota are taxa that may be temporally stable, keystones to their communities, functional to their hosts as facultative symbionts, host-adapted as obligate symbionts, or common across a host population (Risely 2020). Here, we defined the core community as ASVs common to at least 50 % of skin samples of each exposure category (‘before’ or ‘after’) within each treatment group. In total, there were thirty-nine core ASVs in our study. We found the observed ASV richness of the core community was reduced for the ‘classroom’ group after exposure on days two and three (Figure 3a), suggesting minimal recovery in common bacterial associates was possible within a classroom setting (n.b., effective number and Faith’s phylogenetic diversity of ASVs were not significant, Table S3). In contrast, core richness was not different for the ‘sports field’ nor ‘forest’ groups (Figure 3a) after exposure nor across days. However, the compositional change of bacterial communities was strongest in the ‘forest’ group when comparing ‘before’ and ‘after’ exposure samples (R^2^ = 0.33) relative to the ‘sports field’ (R^2^ = 0.26) and ‘classroom’ (R^2^ = 0.24) groups (Figure 3c, Table S5). This indicates that ASV turnover in the ‘forest’ and ‘sports field’ groups’ core skin microbiota buffered any decrease in diversity from the disturbance. We also note that core bacterial community structure on the skin of the ‘classroom’ (R^2^ = 0.01, P > 0.05, Figure 3e, Table S5) and ‘sports field’ (R^2^ = 0.04, P > 0.05) groups did not have significant changes ‘before’ and ‘after’ exposure, although the ‘forest’ group did (R^2^ = 0.07, P < 0.05). In line with our study hypothesis, exposure to green spaces thus enabled the recovery and enrichment of a diverse core community within a short space of time (i.e., 45 mins). Further, exposure to a higher biodiversity setting (i.e., forest) provided a stronger effect compared to a lower biodiversity setting (i.e., sports field or classroom).

**Figure 3.**
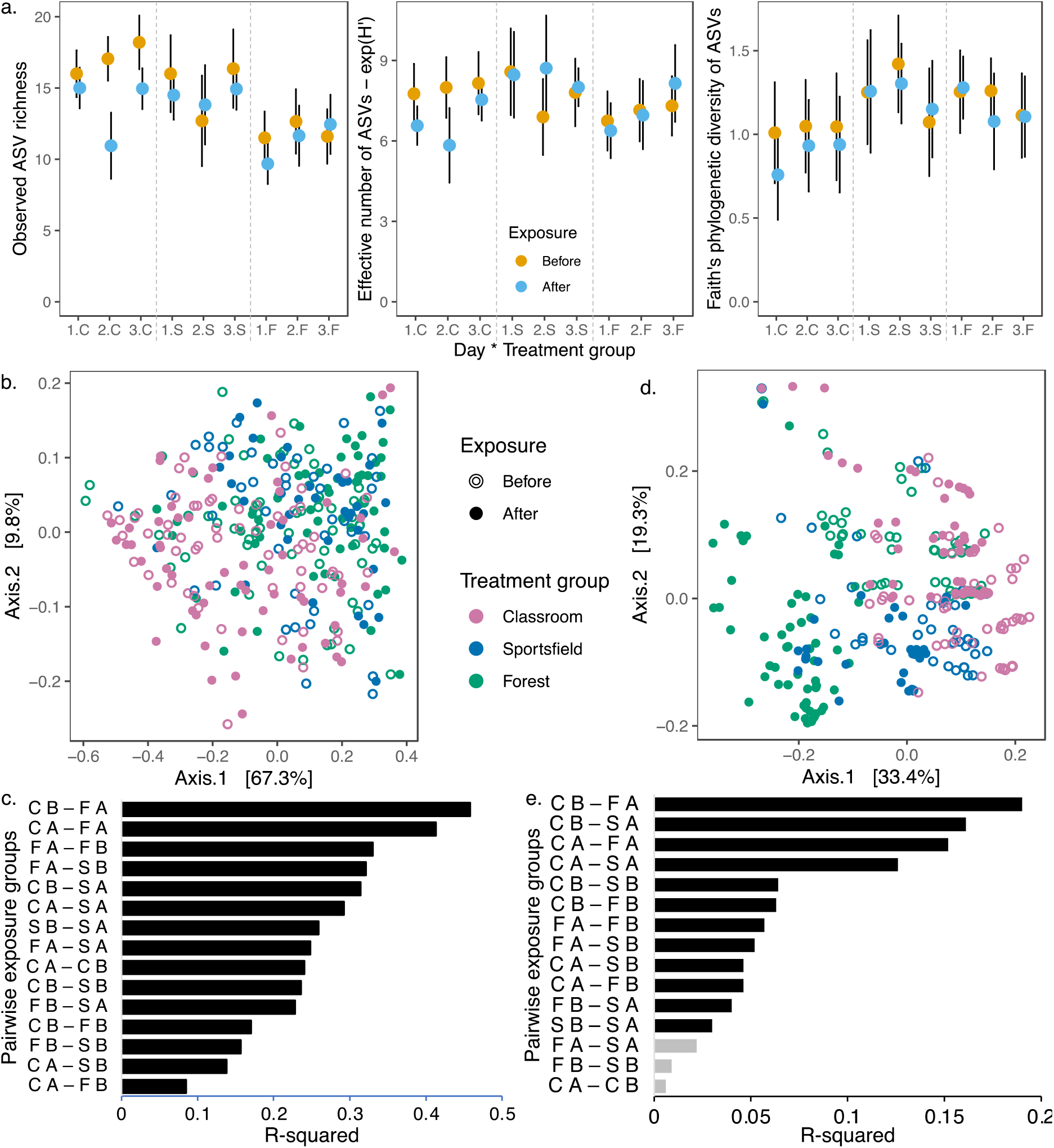
Core bacterial ASV communities of children’s wrists before and after exposure to school yard environments. **a**. Observed richness, effective number (exponent of Shannon’s diversity), and Faith’s phylogenetic diversity of ASVs are shown from the wrists of children exposed to three different school yard environments over three consecutive days. Points are means ± 95 % confidence interval. Significantly different pairs are listed in Table S5. 1, day 1; 2, day 2; 3, day 3; C, ‘classroom’; S, ‘sports field’; F, ‘forest’. **b. & d**. PCoA analyses of weighted-UniFrac and unweighted-UniFrac values, respectively, from all skin samples taken ‘before’ and ‘after’ outdoor exposure. **c. & e**. Bar plots ranking R^2^ values to show strength of difference between pairs of environmental exposure groups (i.e., ‘classroom’ group ‘after’ exposure vs. ‘classroom’ group ‘before’ exposure, CA – CB) from the weighted-UniFrac and unweighted-UniFrac PCoAs, respectively. C, ‘classroom’; S, ‘sports field’; F, ‘forest’; B, ‘before’ exposure; A, ‘after’ exposure. Grey bars represent no significant difference between pairs, black bars represent significant difference where α = 0.05.

**Figure 4.**
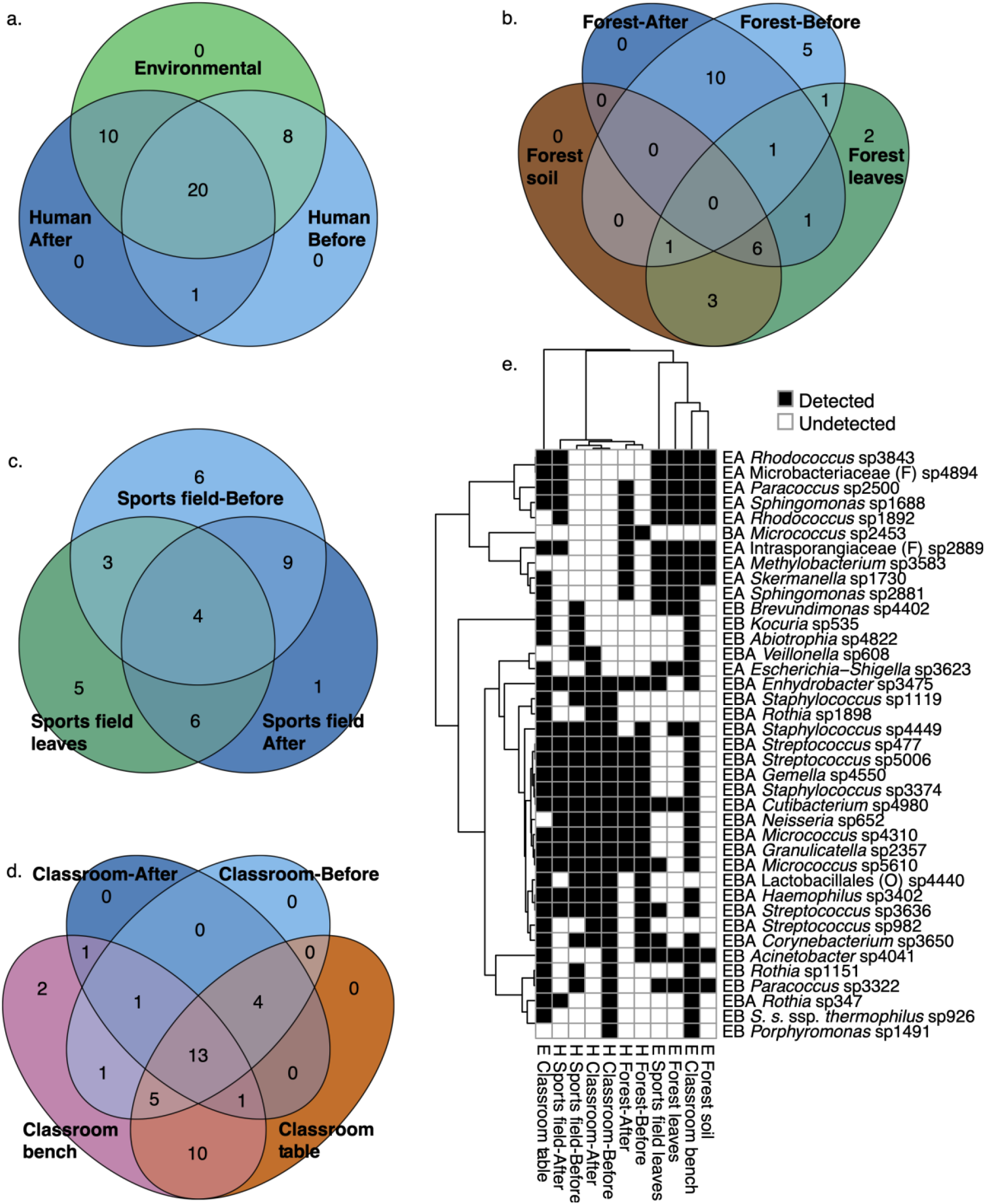
Shared and unshared core bacterial community ASVs between human samples and environmental samples. **a**. Total shared and unshared bacterial ASVs between environmental samples and human samples collected ‘before’ and ‘after’ exposure. **b**. Total shared and unshared bacterial ASVs between the forest environmental samples (soil and leaf surfaces) and human samples from the ‘forest’ treatment group collected ‘before’ and ‘after’ exposure. **c**. Total shared and unshared bacterial ASVs between the ‘sports field’ environmental samples (leaf surfaces) and human samples from the ‘sports field’ treatment group collected ‘before’ and ‘after’ exposure. **d**. Total shared and unshared bacterial ASVs between the ‘classroom’ environmental samples (bench tops and work tables) and human samples from the ‘classroom’ treatment group collected ‘before’ and ‘after’ exposure. **e**. A heatmap of detected community bacterial ASVs by sample type with clustering representing Pearson correlation between columns (samples) and between rows (ASVs). H and E on the x-axis represent human and environmental sample types, respectively. Y-axis lettering represents detection in environmental (E), human-before exposure (B), and/or human-after exposure (A) samples. All ASVs are named by their lowest identified taxonomic rank, and *S. s*. ssp. *thermophilus* is *Streptococcus salivarius* ssp. *thermophilus*.

Lastly, we investigated the community turnover and transfer of new ASVs into the core microbiota that occurred through green space exposure. We first identified which bacterial ASVs were lost from the core community. Of the original 29 core ASVs across the three groups, eight were lost, while ten were gained from the exposure to treatment environments (Figure 3a). In the ‘forest’ group, seven core ASVs were gained from the environment (six were found in forest soil and on leaves; one on forest leaves), after losing seven from pre-disturbance (Figure 3b). The ‘sports field’ group gained seven new core ASVs (six were found on sports field leaves; one of unknown origin), after losing nine (Figure 3c). The ‘classroom’ group lost six core ASVs, and only gained two (Figure 3d) (one found on classroom benchtops and one on tables, with one of these belonging to the potentially pathogenic group *Escherichia*-*Shigella* (Peng et al. 2009; Figure 3e). The potential health effect of the higher core ASV turnover on the skin microbiota for the ‘forest’ treatment group remains unclear.

While the long-term health impacts of the rapid changes in the skin microbiota seen here require futher study, a diverse microbiota has often been correlated to positive health outcomes (Hanski et al. 2012; Stein et al. 2016; Birzele et al. 2017; Egert et al. 2017; Roslund et al. 2020). Indeed, our findings have implications for the quality and diversity of plantings to be used in urban green spaces, especially in school grounds. The quality of biodiversity in our environments is most likely what provides the human microbiota with enrichment of diverse microbes and resilience to disturbance – the ability to maintain diversity in a dynamic ecology between environment, host, and microorganisms (Shade et al. 2012). However, the factors and mechanisms that underpin this environmentally enabled resilience and how this may be related to long-term health outcomes require further investigation. Nevertheless, we are confident that biodiversity interventions of urban green space will have a positive effect on public health that can transcend socio-economic boundaries for health care (Mills et al. 2019). It is also important to note that if measures are not taken to prevent ‘green gentrification’ – the increasing exclusivity of urban greening linked to socioeconomic status – then biodiversity interventions will not help those that are the most in need of cost-effective primary health preventions (Jelks et al. 2021). The first place to start ensuring that people receive adequate access to diverse environmental microorganisms is in schools.

## Methods

### Ethics

This project was done under ethics approval by The University of Adelaide’s Human Research Ethics Committee (approval number H-2019-064) and by the South Australian Government’s Department for Education (approval number 2019-7388569).

### Metadata

Prior to participating in sampling, the school students and their parents or guardians were asked to complete a questionnaire about exclusion criteria: antibiotic use in the previous six-months, allergies to sampling materials, and skin conditions.

### Design and sampling

The study site was a primary school in Adelaide, Kaurna Country, South Australia, Australia. Participants were 10- to 11-year-old children, who were students of the school. We requested consent for 90 participants with an uptake rate of 80% (*n* = 72). Fifty-seven students (*n* female = 24) participants passed our exclusion criteria.

The participants were already divided into three classes and had spent approximately nine months of the 2019 school year in this configuration before sampling. We assigned the classes to one of three treatment groups, or schoolyard locations – ‘classroom’, ‘sports field’, or ‘forest’ – based on their teachers pre-existing proclivity to spend time in these environments to not disrupt their normal scheduling. We then followed a ‘before’ and ‘after’ exposure sampling regime, where the ‘before’ sampling also acted as a disturbance event to the skin microbiota and therefore provided pre-disturbance information. Participants were swabbed before and after a forty-five-minute exposure to their assigned treatment locations. The combinations of the treatment groups with the exposure created the variable of exposure groups (i.e., treatment group level + exposure level, e.g., ‘classroom before’, ‘classroom after’).

The experiment was repeated three times over three days, 13^th^ November to 15^th^ November 2019, where we ran 1-hour sampling sessions from 9am to 10am each day. After spending approximately 30-60 minutes in the classroom, the sampling sessions started with a ‘before’ sample in the classroom, followed by 45-minutes of standard school activities in their treatment locations followed by an ‘after’ sample back in the classroom. The ‘before’ exposure samples on day one allowed us to test for differences in long-term microbiota divergence between groups over the course of the 2019 school year.

Samples were a skin swab collected by applying 2 drops of sterile saline solution (Reclens Saline Solution, Aaxis Pacific, Blacktown, Australia) to the inside of the participant’s left wrist (as in Selway et al. 2020) followed by having them rub a nylon FLOQ swab (COPAN, Brescia, Italy) according to manufacturer’s instructions in an area 3 cm in diameter. The environments were also sampled for comparison to the human samples with swabs taken of the classroom tables (desks) and benches (sideboards) (*n* = 8), the sports field grass (*n* = 6), and the forest leaf-surfaces (*n* = 4) and soil (*n* = 4). The swab tips of human and environmental samples were collected in 1 mL eNAT DNA stabilisation solution (COPAN, Brescia, Italy) and stored at -20 ºC until DNA extraction.

### DNA Extraction, PCR, Library Preparation, and Bioinformatics

DNA extractions were done across two different laboratories with different technicians due to COVID-19 restrictions. Samples were randomly assigned to the extraction labs. The first lab, Australian DNA Identification and Forensic Facility (ADIFF), randomly selected one sample from each sampling group (i.e., ‘treatment group’ x ‘exposure’ x ‘day’, or environmental sample) for each extraction batch done in that lab (*n* samples extracted at ADIFF = 140) and the remainder (*n* samples = 202) were sent to the second lab, the Australian Genome Research Facility (AGRF). DNA was extracted from human and environmental samples in both laboratories using DNeasy PowerSoil Pro Kit (QIAGEN) as per the manufacturer’s instructions. Extraction blank controls (EBCs) were used for all ADIFF processed samples.

PCR amplification and sequencing of all samples were done by the AGRF. Bacterial 16S V3-V4 PCR amplicons were generated using the primers and conditions outlined in Table S1. Thermocycling was completed with an Applied Biosystem 384 Veriti and using Platinum SuperFi II master mix (Invitrogen, Australia) for the primary PCR. The first stage PCR was cleaned using magnetic beads (Beckman Coulter, SPRI), and samples were visualised on 2 % Sybr Egel (Thermo-Fisher). A secondary PCR to index the amplicons was performed with the same polymerase master mix. The resulting amplicons were cleaned again using magnetic beads, quantified by fluorometry (Promega Quantifluor) and normalised. The equimolar pool of all amplicons was cleaned a final time using magnetic beads to concentrate the pool and then measured using a High-Sensitivity D1000 Tape on an Agilent 2200 TapeStation. The pool was diluted to 5 nM and molarity was confirmed again using a Qubit High Sensitivity dsDNA assay (ThermoFisher). This was followed by sequencing on an Illumina MiSeq (San Diego, CA, USA) with a V3, 600 cycle kit (2 x 300 base pairs paired-end).

### Bioinformatics

Pre-processing was done by AGRF using QIIME2 (Bolyen et al. 2019) version 2019.7. Samples were demultiplexed using Illumina scripts. Raw sequences were searched and trimmed for template-specific primers using Cutadapt with default quality settings (Martin 2011). Amplicon sequence variants (ASVs) were then generated at 240 bases using DADA2 (Callahan et al. 2016). Taxonomy was assigned to ASVs with the Silva 132 ‘sklearn’ classifier using a trained database for the 16S V3-V4 gene region (Quast et al. 2012).

We removed ASVs that were 100 % biased to one extraction lab or the other. Further, we identified contaminant ASVs from non-template EBC and PCR controls using the prevalence method within the *decontam* package (v 1.8.0; Davis et al. 2018) in R (v 4.0.0; RCoreTeam 2019) and with a threshold probability of 0.5. Any identified contaminants were removed from all biological samples before downstream analysis. Additionally, ASVs assigned to mitochondria, chloroplast, Archaea, or ‘unknown’ were removed, and ASVs found in fewer than two biological samples in the dataset or with fewer than 9 reads (Edgar 2016) across all samples were also excluded. After pre-processing there were 5,412 ASVs with reference sequences and a total 18,835,659 sequences across 342 samples. Unrooted phylogenetic trees were constructed using the *msa* package version 1.16.0 (Bodenhofer et al. 2015) for multiple sequence alignment and the *phangorn* package version 2.5.5 (Schliep 2010) for phylogenetic tree building.

### Core-community of human skin samples

Due to the inherent variability between individuals, we determined core bacterial communities to test for experimental changes to the wrist community. To determine the core community, we divided the main dataset of human skin samples into six subsets based on the six exposure groups (i.e., treatment group by exposure combinations) – ‘classroom before’, ‘classroom after’, ‘sports field before’, ‘sports field after’, ‘forest before’, and ‘forest after’ – with the ‘subset_samples’ function of the *phyloseq* package (v 1.32.1; McMurdie & Holmes 2013). We then used the ‘ps_prune’ function of the *MicEco* package to keep only those ASVs that were present in at least 50 % of the samples within each of these subsets. Once the ≥ 50 % prevalent ASVs were identified, they were merged back into a single dataset using the ‘merge_phyloseq’ function of the *phyloseq* package. This process identified 39 ASVs as core to the skin samples of this project at ≥ 50 % prevalence. We constructed an unrooted phylogenetic tree for the core community as above. We then merged the data of those 39 ASVs from the environmental samples into the human core community dataset for comparison between human and environmental sample types. From the 39 core ASVs, there were 6,451,217 total sequences across human and environmental samples.

### Statistics

All statistics were calculated in R (v 4.0.0; RCoreTeam 2019). Three datasets were analysed, human skin communities (from skin swabs), core human skin communities (from skin swabs), and environmental communities (from soil, leaf surface, and classroom surface swabs).

Before alpha diversity was calculated, the filtered ASV datasets were rarefied to 3,124 reads for the human skin communities, 1,103 reads for the core human skin communities, and 10,933 reads for the environmental communities with the ‘rarefy_even_depth’ function of the *phyloseq* package (v 1.32.1; McMurdie & Holmes 2013). Alpha diversity was calculated as observed ASV richness and Shannon’s diversity with the ‘estimate_richness’ function in *phyloseq* and Faith’s phylogenetic diversity was calculated with the ‘pd’ function of the *picante* package (v 1.8.1; Kembel et al. 2010). We converted Shannon’s diversity to effective number of ASVs by taking its exponent (Jost 2006). We used generalized linear mixed models (GLMMs) to test for difference in alpha diversity by crossing the fixed factors of ‘treatment group’, ‘exposure’, and ‘day’ and adding the random factors of ‘student id’, ‘student id interacting with day’ to account for repeat sampling, and ‘student id interacting with exposure’ to account for repeat exposures. GLMMs were done with the ‘glmer’ function of the *lme4* package v 1.1-25 (v 1.1-25; Bates et al. 2007). Distributions for the GLMMs were negative-binomial for observed ASV richness (count data) and Gamma for Faith’s phylogenetic diversity and effective number of ASVs (Shannon’s), which were positive, non-integer, and non-parametric. Main effects of the GLMMs were tested by Type II Wald Chi^2^ tests with the ‘Anova’ function of the *car* package (v 3.0-10; Fox et al. 2012). Pairwise comparisons of ‘treatment group’, ‘exposure’, and ‘day’ combinations were done by z-tests with Tukey P-value adjustment with the ‘emmeans’ function of the *emmeans* package (v 1.6.0; Lenth 2018).

Ordinations of beta diversity on the three datasets were done with the ‘ordinate’ function in *phyloseq*. Ordinations were based on unrarefied data in principal coordinates analysis (PCoA) with weighted-UniFrac and unweighted-UniFrac distance matrices. We used PERMANOVA, with 999 iterations, with the ‘adonis’ function of the *vegan* package (v 2.5-6; Oksanen et al. 2017) to test the model of ‘treatment group’ by ‘exposure’ by ‘day’. Pairwise comparisons between exposure groups (e.g. ‘forest before’ vs. ‘forest after’) were tested by PERMANOVA with 999 iterations with the ‘pairwise.adonis2’ function of the *pairwise*.*adonis* package (v 0.0.1; Arbizu 2017).

Shared ASVs between environmental, ‘before’ exposure, and ‘after’ exposure human samples were tallied using the ‘ps_venn’ function of the *MicEco* package (v 0.9.15; Russel 2021). ASVs were plotted by sample type into detected/undetected heatmaps using the ‘pheatmap’ function of the *pheatmap* package (v 1.0.12; Kolde & Kolde 2015). The heatmap cells were clustered based on Pearson correlation between rows and columns.

### Data access

Raw sequence data is stored on the Sequence Read Archive server with the BioProject ID: PRJNA738964. Find the Phyloseq compatible metadata, ASV table, taxonomy table, ASV reference sequences, and R script used in this analysis, along with relevant ethics approvals on Figshare with the DOI: 10.25909/14787867.

## Acknowledgments and declarations

Ethics approval was given by The University of Adelaide’s Human Research Ethics Committee (H-2019-064) and by The Government of South Australia’s Department for Education (2019-7388569). Written consent for the participants was given by parents or caregivers. All interactions with participants, from consent to sampling and data handling, was done according to the National Statement on Ethical Conduct of Human Research (2007) and The Australian Code for the Responsible Conduct of Research (2018). This project was funded by The University of Adelaide. Jacob Mills and Caitlin Selway were studying under Australian Commonwealth Research Training Program Stipends. We declare no conflict of interest.

We would like to express our deep gratitude to the school staff for opening their doors to us and being open to our research, and for working hard and collaboratively with us on this project. We would also like to thank the students and parents for their participation in this study. We thank Philip Weinstein and Patrick O’Connor for assistance in initiating the study. We would like to thank the hard working volunteers, Helen Mills, James Loader, and Ana Judith Giraldo, who lead the teachers and participants during the sampling sessions and oversaw the scientific integrity of the sampling. We thank the staff at AGRF, especially Naga Rup Pinaki Kasinadhuni, for providing efficient, flexible and ongoing service for our bioinformatics needs. The authors acknowledge Martin Breed for his contributions.

