## Supplementary Material for "School yard biodiversity determines short-term recovery of disturbed skin microbiota in children"

**Table S1** PCR program and primers at the Australian Genome Research Facility.

| Target | Cycle | Initial | Disassociate | Anneal | Extension | Finish |
|---|---|---|---|---|---|---|
| 16S: V3 - V4 | 29 | 95ºC for 7 min | 94ºC for 30s | 50ºC for 60S | 72ºC for 60S | 72ºC for 7 min |

| Target | 341F-806R |
|---|---|
| Forward Primer (341F) | CCTAYGGGRBGCASCAG |
| Reverse Primer (806R) | GGACTACNNGGGTATCTAAT |

**Table S2 Alpha diversity of bacterial (16S rRNA ASV) communities of student's wrists before and after exposure to either forest, sports field, or classroom environments.** EN is effective number of ASVs calculated as the exponent of Shannon's diversity index. Only showing significantly different pairs for at least one diversity metric. Significance codes Pr(>$Chi^2$): 'ns' not significant; 'º' P < 0.10; '*' P < 0.05; '**' P < 0.01; '***' P < 0.001.

| GLMM - Type II Wald $Chi^2$ test ~ Treatment * Exposure * Day | Observed ASV richness | | | EN (Shannon's) | | | Faith's PD | | |
|---|---|---|---|---|---|---|---|---|---|
| | $Chi^2$ | Pr(>$Chi^2$) | Sig. | $Chi^2$ | Pr(>$Chi^2$) | Sig. | $Chi^2$ | Pr(>$Chi^2$) | Sig. |
| Treatment group | 0.08 | 0.963 | ns | 1.51 | 0.470 | ns | 1.55 | 0.462 | ns |
| Exposure | 1.58 | 0.209 | ns | 10.25 | 0.001 | ** | 10.69 | 0.001 | ** |
| Day | 9.95 | 0.007 | ** | 4.18 | 0.124 | ns | 1.88 | 0.391 | ns |
| Treatment * Exposure | 8.62 | 0.013 | * | 3.97 | 0.137 | ns | 2.99 | 0.224 | ns |
| Treatment * Day | 27.52 | < 0.001 | *** | 11.09 | 0.026 | * | 7.21 | 0.125 | ns |
| Exposure * Day | 0.62 | 0.734 | ns | 1.12 | 0.573 | ns | 1.37 | 0.503 | ns |
| Treatment * Exposure * Day | 16.00 | 0.003 | ** | 10.61 | 0.031 | * | 17.85 | 0.001 | ** |
| **Pairwise GLMM ~ Treatment * Exposure * Day** | **Observed ASV richness** | | | **EN (Shannon's)** | | | **Faith's PD** | | |
| | z-ratio | P | Sig. | z-ratio | P | Sig. | z-ratio | P | Sig. |
| Classroom Before 1 – Forest After 2 | -2.47 | 0.551 | ns | 2.99 | 0.203 | ns | 3.78 | 0.018 | * |
| Forest Before 1 – Forest After 2 | -4.23 | 0.003 | ** | 3.53 | 0.044 | * | 3.42 | 0.063 | º |
| Forest Before 1 – Forest After 3 | -3.88 | 0.013 | * | 3.29 | 0.092 | º | 2.58 | 0.469 | ns |
| Forest After 1 – Forest After 2 | -4.27 | 0.003 | ** | 3.33 | 0.082 | º | 3.35 | 0.076 | º |
| Forest After 1 – Forest After 3 | -3.93 | 0.011 | * | 3.08 | 0.162 | ns | 2.38 | 0.623 | ns |
| Sports field Before 2 – Forest After 2 | -2.99 | 0.205 | ns | 2.86 | 0.278 | ns | 3.91 | 0.011 | * |
| Forest Before 2 – Forest After 2 | -2.00 | 0.869 | ns | 2.94 | 0.230 | ns | 4.03 | 0.007 | ** |
| Classroom After 2 – Forest After 2 | -3.99 | 0.008 | ** | 3.55 | 0.041 | * | 3.14 | 0.141 | ns |
| Classroom After 2 – Classroom Before 3 | -4.23 | 0.003 | ** | 1.90 | 0.911 | ns | 0.88 | 1.000 | ns |
| Classroom After 2 – Forest After 3 | -3.68 | 0.026 | * | 3.35 | 0.078 | º | 2.34 | 0.651 | ns |
| Forest Before 3 – Forest After 3 | -3.14 | 0.140 | ns | 3.56 | 0.040 | * | 2.93 | 0.237 | ns |

**Table S3 Alpha diversity of core bacterial (16S rRNA ASV) communities of student's wrists before and after exposure to either forest, sports field, or classroom environments.** EN is effective number of ASVs calculated as the exponent of Shannon's diversity index. Only showing significantly different pairs for at least one diversity metric. Significance codes Pr(>$Chi^2$): 'ns' not significant; 'º' P < 0.10; '*' P < 0.05; '**' P < 0.01; '***' P < 0.001.

| GLMM - Type II Wald $Chi^2$ test ~ Treatment * Exposure * Day | Observed ASV richness | | | EN (Shannon's) | | | Faith's PD | | |
|---|---|---|---|---|---|---|---|---|---|
| | $Chi^2$ | Pr(>$Chi^2$) | Sig. | $Chi^2$ | Pr(>$Chi^2$) | Sig. | $Chi^2$ | Pr(>$Chi^2$) | Sig. |
| Treatment group | 27.00 | <0.001 | *** | 1.07 | 0.587 | ns | 1.85 | 0.396 | ns |
| Exposure | 15.62 | <0.001 | *** | 0.77 | 0.381 | ns | 0.21 | 0.648 | ns |
| Day | 7.59 | 0.022 | * | 2.92 | 0.232 | ns | 0.16 | 0.923 | ns |
| Treatment * Exposure | 7.85 | 0.020 | * | 5.60 | 0.061 | º | 0.25 | 0.883 | ns |
| Treatment * Day | 10.56 | 0.032 | * | 3.04 | 0.551 | ns | 0.50 | 0.974 | ns |
| Exposure * Day | 1.70 | 0.427 | ns | 3.39 | 0.183 | ns | 0.08 | 0.962 | ns |
| Treatment * Exposure * Day | 14.42 | 0.006 | ** | 7.40 | 0.116 | ns | 0.19 | 0.996 | ns |

| Pairwise GLMM ~ Treatment * Exposure * Day | Observed ASV richness | | | EN (Shannon's) | | | Faith's PD | | |
|---|---|---|---|---|---|---|---|---|---|
| | z-ratio | P | Sig. | z-ratio | P | Sig. | z-ratio | P | Sig. |
| Classroom Before 1 – Forest Before 1 | 3.50 | 0.049 | * | -1.17 | 1.000 | ns | 0.50 | 1.000 | ns |
| Classroom Before 1 – Forest After 1 | 5.09 | < 0.001 | *** | -1.55 | 0.987 | ns | 0.54 | 1.000 | ns |
| Classroom Before 1 – Classroom After 2 | 4.21 | 0.003 | ** | -2.45 | 0.566 | ns | -0.18 | 1.000 | ns |
| Sports field Before 1 – Forest After 1 | 4.63 | < 0.001 | *** | -1.73 | 0.962 | ns | 0.05 | 1.000 | ns |
| Forest Before 1 – Classroom Before 2 | -4.20 | 0.004 | ** | 1.13 | 1.000 | ns | -0.42 | 1.000 | ns |
| Forest Before 1 – Classroom Before 3 | -4.94 | < 0.001 | *** | 1.52 | 0.990 | ns | -0.42 | 1.000 | ns |
| Classroom After 1 – Forest After 1 | 4.39 | 0.002 | ** | -0.45 | 1.000 | ns | 1.15 | 1.000 | ns |
| Sports field After 1 – Forest After 1 | 3.52 | 0.045 | * | -1.72 | 0.964 | ns | 0.04 | 1.000 | ns |
| Forest After 1 – Classroom Before 2 | -5.77 | < 0.001 | *** | 1.51 | 0.990 | ns | -0.46 | 1.000 | ns |
| Forest After 1 – Classroom Before 3 | -6.50 | < 0.001 | *** | 1.89 | 0.917 | ns | -0.47 | 1.000 | ns |
| Forest After 1 – Sports field Before 3 | -4.85 | < 0.001 | *** | 1.40 | 0.996 | ns | -0.36 | 1.000 | ns |
| Forest After 1 – Classroom After 3 | -4.36 | 0.002 | ** | 1.51 | 0.990 | ns | -0.71 | 1.000 | ns |
| Forest After 1 – Sports field After 3 | -3.94 | 0.010 | * | 1.67 | 0.972 | ns | -0.22 | 1.000 | ns |
| Classroom Before 2 – Classroom After 2 | 4.96 | < 0.001 | *** | -3.16 | 0.133 | ns | -0.26 | 1.000 | ns |
| Classroom Before 2 – Forest After 2 | 3.92 | 0.011 | * | -0.66 | 1.000 | ns | 0.06 | 1.000 | ns |
| Classroom Before 2 – Forest Before 3 | 3.97 | 0.009 | ** | -0.51 | 1.000 | ns | 0.14 | 1.000 | ns |
| Forest Before 2 – Classroom Before 3 | -3.85 | 0.014 | * | 0.83 | 1.000 | ns | -0.42 | 1.000 | ns |
| Classroom After 2 – Classroom Before 3 | -5.76 | < 0.001 | *** | 2.85 | 0.279 | ns | 0.25 | 1.000 | ns |
| Classroom After 2 – Sports field Before 3 | -3.60 | 0.035 | * | 1.88 | 0.920 | ns | 0.29 | 1.000 | ns |
| Forest After 2 – Classroom Before 3 | -4.64 | < 0.001 | *** | 1.04 | 1.000 | ns | -0.07 | 1.000 | ns |
| Classroom Before 3 – Forest Before 3 | 4.69 | < 0.001 | *** | -0.90 | 1.000 | ns | 0.14 | 1.000 | ns |
| Classroom Before 3 – Forest After 3 | 4.02 | 0.007 | ** | -0.32 | 1.000 | ns | 0.13 | 1.000 | ns |

**Table S4** Main (with homogeneity of dispersion tests, Disp.) and pairwise PERMANOVA on bacterial ASV communities of student's wrists before and after exposure to assigned school environments. Significance codes Pr(>F): 'ns' not significant; '°' P < 0.10; '*' P < 0.05; '**' P < 0.01; '***' P < 0.001.

| Main PERMANOVA | | Weighted-UniFrac | | | | Unweighted-UniFrac | | | |
|---|---|---|---|---|---|---|---|---|---|
| distance ~ Treatment*Exposure*Day | | $R^2$ | F | Pr(>F) | Disp. | $R^2$ | F | Pr(>F) | Disp. |
| Treatment | $df_{2,321}$ | 0.09 | 16.29 | *** | ** | 0.02 | 3.94 | *** | *** |
| Exposure | $df_{1,321}$ | 0.02 | 7.09 | *** | ns | 0.01 | 2.29 | *** | ** |
| Day | $df_{2,321}$ | 0.01 | 2.02 | * | ns | 0.01 | 1.57 | ** | ** |
| Treatment*Exposure | $df_{2,321}$ | 0.01 | 1.94 | * | ° | 0.01 | 1.65 | ** | ° |
| Treatment*Day | $df_{4,321}$ | 0.01 | 1.11 | ns | | 0.02 | 1.36 | ** | |
| Exposure*Day | $df_{2,321}$ | 0.01 | 1.00 | ns | | 0.01 | 1.05 | ns | |
| Treatment*Exposure*Day | $df_{4,321}$ | 0.01 | 0.98 | ns | | 0.01 | 1.22 | * | ** |
| **Pairwise PERMANOVA** | | Weighted-UniFrac | | | | Unweighted-UniFrac | | | |
| Distance ~ Treatment*Exposure | | $R^2$ | F | Pr(>F) | | $R^2$ | F | Pr(>F) | |
| Forest Before - Classroom Before | $df_{1,121}$ | 0.06 | 8.14 | *** | | 0.02 | 2.44 | *** | |
| Forest Before - Sports field Before | $df_{1,102}$ | 0.01 | 0.96 | ns | | 0.02 | 1.70 | * | |
| Classroom Before - Sports field Before | $df_{1,100}$ | 0.06 | 6.82 | *** | | 0.01 | 1.48 | * | |
| Forest After – Forest Before | $df_{1,123}$ | 0.06 | 7.43 | *** | | 0.02 | 2.95 | *** | |
| Sports field After - Sports field Before | $df_{1,77}$ | 0.03 | 2.37 | * | | 0.02 | 1.52 | * | |
| Classroom After - Classroom Before | $df_{1,119}$ | 0.01 | 0.72 | ns | | 0.01 | 0.98 | ns | |
| Forest After – Classroom After | $df_{1,121}$ | 0.15 | 21.49 | *** | | 0.04 | 5.11 | *** | |
| Forest After – Sports field After | $df_{1,98}$ | 0.02 | 2.17 | ° | | 0.03 | 2.72 | *** | |
| Classroom After - Sports field After | $df_{1,96}$ | 0.13 | 13.65 | *** | | 0.02 | 2.38 | *** | |
| Forest After – Classroom Before | $df_{1,121}$ | 0.19 | 28.20 | *** | | 0.05 | 6.03 | *** | |
| Forest After – Sports field Before | $df_{1,102}$ | 0.05 | 5.49 | *** | | 0.03 | 3.64 | *** | |
| Classroom After - Forest Before | $df_{1,121}$ | 0.05 | 5.77 | *** | | 0.02 | 1.99 | *** | |
| Classroom After - Sports field Before | $df_{1,100}$ | 0.05 | 4.76 | ** | | 0.01 | 1.43 | * | |
| Sports field After - Forest Before | $df_{1.98}$ | 0.04 | 4.04 | ** | | 0.02 | 1.61 | * | |
| Sports field After - Classroom Before | $df_{1,96}$ | 0.16 | 18.18 | *** | | 0.03 | 2.58 | *** | |

**Table S5** Main (with homogeneity of dispersion tests, Disp.) and pairwise PERMANOVA on core bacterial ASV communities of student's wrists before and after exposure to assigned school environments. Significance codes Pr(>F): 'ns' not significant; 'º' P < 0.10; '*' P < 0.05; '**' P < 0.01; '***' P < 0.001.

| Main PERMANOVA | | Weighted-UniFrac | | | | Unweighted-UniFrac | | | |
|---|---|---|---|---|---|---|---|---|---|
| distance ~ Treatment*Exposure*Day | | $R^2$ | F | Pr(>F) | Disp. | $R^2$ | F | Pr(>F) | Disp. |
| Treatment | $df_{2,321}$ | 0.11 | 19.59 | *** | º | 0.15 | 38.59 | *** | ** |
| Exposure | $df_{1,321}$ | 0.02 | 7.04 | ** | ns | 0.10 | 51.25 | *** | * |
| Day | $df_{2,321}$ | 0.01 | 1.18 | ns | ns | 0.01 | 1.58 | º | º |
| Treatment*Exposure | $df_{2,321}$ | 0.02 | 3.03 | * | º | 0.15 | 38.46 | *** | ns |
| Treatment*Day | $df_{4,321}$ | 0.01 | 0.63 | ns | | 0.01 | 1.34 | ns | |
| Exposure*Day | $df_{2,321}$ | <0.01 | 0.86 | ns | | 0.01 | 1.42 | ns | |
| Treatment*Exposure*Day | $df_{4,321}$ | <0.01 | 0.40 | ns | | 0.01 | 1.06 | ns | |
| Pairwise PERMANOVA | | Weighted-UniFrac | | | | Unweighted-UniFrac | | | |
| Distance ~ Treatment*Exposure | | $R^2$ | F | Pr(>F) | | $R^2$ | F | Pr(>F) | |
| Forest Before - Classroom Before | $df_{1,121}$ | 0.06 | 8.22 | *** | | 0.17 | 24.51 | *** | |
| Forest Before - Sports field Before | $df_{1,102}$ | 0.01 | 1.35 | ns | | 0.16 | 18.85 | *** | |
| Classroom Before - Sports field Before | $df_{1,100}$ | 0.09 | 9.50 | *** | | 0.24 | 30.62 | *** | |
| Forest After – Forest Before | $df_{1,123}$ | 0.07 | 9.84 | *** | | 0.33 | 60.21 | *** | |
| Sports field After - Sports field Before | $df_{1,77}$ | 0.04 | 2.78 | º | | 0.26 | 26.58 | *** | |
| Classroom After - Classroom Before | $df_{1,119}$ | 0.01 | 1.27 | ns | | 0.24 | 37.12 | *** | |
| Forest After – Classroom After | $df_{1,121}$ | 0.18 | 26.10 | *** | | 0.41 | 84.34 | *** | |
| Forest After – Sports field After | $df_{1,98}$ | 0.02 | 1.97 | ns | | 0.25 | 31.91 | *** | |
| Classroom After - Sports field After | $df_{1,96}$ | 0.15 | 16.37 | *** | | 0.29 | 39.17 | *** | |
| Forest After – Classroom Before | $df_{1,121}$ | 0.20 | 30.53 | *** | | 0.46 | 101.4 | *** | |
| Forest After – Sports field Before | $df_{1,102}$ | 0.05 | 5.81 | ** | | 0.32 | 47.78 | *** | |
| Classroom After - Forest Before | $df_{1,121}$ | 0.06 | 7.32 | ** | | 0.08 | 11.19 | *** | |
| Classroom After - Sports field Before | $df_{1,100}$ | 0.08 | 8.24 | ** | | 0.14 | 15.85 | *** | |
| Sports field After - Forest Before | $df_{1.98}$ | 0.05 | 4.96 | ** | | 0.23 | 28.71 | *** | |
| Sports field After - Classroom Before | $df_{1,96}$ | 0.17 | 19.62 | *** | | 0.31 | 43.51 | *** | |

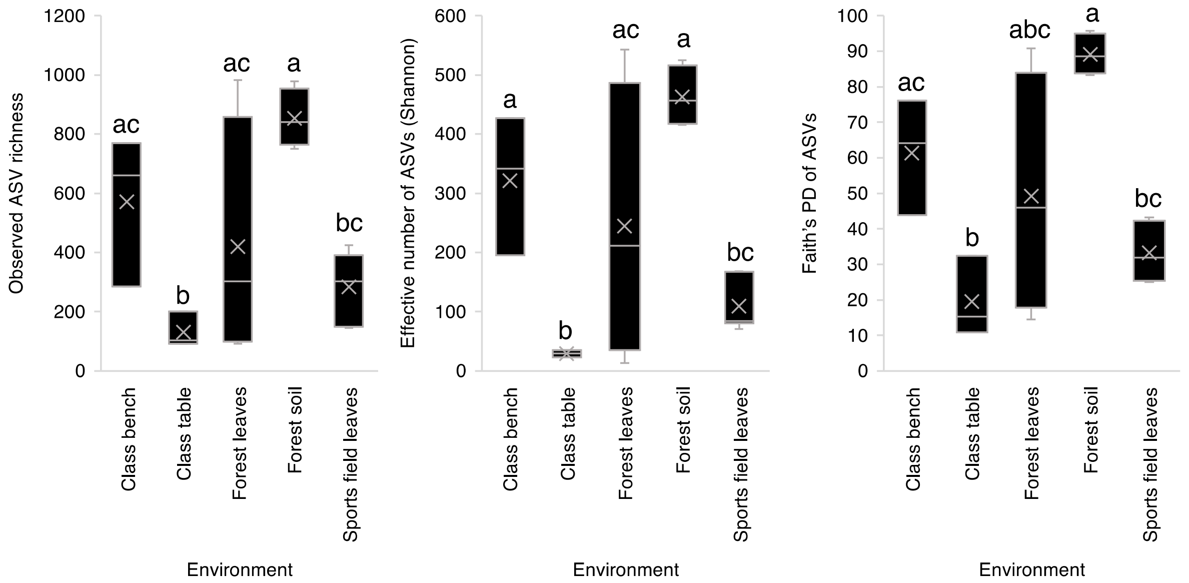


**Figure S1** Alpha diversity of environmental samples from the exposure environments. Shared letters indicate no significant difference (P > 0.05) between pairs.
